# ecode: An R package to investigate community dynamics in ordinary differential equation systems

**DOI:** 10.1101/2023.06.23.546319

**Authors:** Haoran Wu

## Abstract

Population dynamical modelling plays a crucial role in understanding ecological populations and making informed decisions for environmental management. However, existing software packages for dynamical system modelling often lack comprehensive integration of techniques and guidelines, limiting their practical usability. This paper introduces ecode, a novel package for modelling ecological populations and communities using ordinary differential equation systems, designed with a user-friendly framework. By following a three-cycle procedure, users can easily construct ecological models and explore their behaviours through a wide range of graphical, analytical, and numerical techniques. The package incorporates advanced techniques such as grid search methods and simulated annealing algorithms, enabling users to iteratively refine their models and achieve accurate predictions. Notably, ecode minimises external dependencies, ensuring robustness and reducing the risk of package failure caused by updates in dependencies. Overall, ecode serves as a valuable tool for ecological modelling, facilitating the exploration of complex ecological systems and the generation of informed predictions and management recommendations.

## Introduction

Population dynamical modelling is often involved in the study of ecological populations and communities (Gurney and Nisbet 1998; Quinn 2003; Buckland et al. 2007; Barbier et al.2018). The structure of ecological communities are often described by a set of differential equations, which allow scientists to link community patterns and various processes such as local demographics (Bonsall and Hastings 2004; Gouhier et al. 2010), species coexistence (Ives 1988; Rafique and Qader 2015), local adaptation potential (Gurney et al. 1992; Cao et al. 2020), and the way species response to changing climate (Urban et al. 2016; Pilowsky et al. 2022). While these models and the theories derived from them have the capability to predict the future outcomes and assess the risk of extinction (Holmes et al. 2007; Ladle 2009; Howell et al. 2020), how to use them efficiently for environmental management poses a significant challenge (Yates et al. 2018; Schuwirth et al. 2019). To ensure effectiveness, accurate predictions rely on appropriate mathematical models (Jørgensen and Bendoricchio 2001; Grimm et al. 2014), adaptability to diverse management scenarios (Cartwright et al. 2016; Getz et al. 2018; Schuwirth et al. 2019) and anthropogenic climate change (Buckland et al. 2007; Thuiller 2007), and the ability to quantify uncertainties (Chowell 2017). Software packages specifically developed for dynamic systems modelling provide user-friendly interfaces for model construction, simulations, and accurate predictions (Costanza and Voinov 2001).

The R language has emerged as a leading platform among ecologists due to its extensive array of ecological analysis toolkits, making it highly favoured within the ecological research community (Zuur et al. 2009; Borcard et al. 2011; Swenson 2014). In the field of dynamical system modelling, a multitude of R packages have been developed, covering a wide range of analytical and numerical methods. The package deSolve is used to deal with initial value problems in differential equation systems (Soetaert et al. 2010). It also serves as an essential dependency for various modelling packages such as phaseR (Grayling 2014), which allows users to investigate differential equations through phase plane methods. Package bvpSolve provides functions to solve boundary value problems (Mazzia et al., 2014). Package fitode provides functions for fitting differential equations using Markov chain Monte Carlo method (Park and Bolker 2022). More intricate differential equations, such as stochastic and delayed differential equations, can be solved by the sde (Iacus 2007; Iacus 2008) and PBSddesolve (Couture-Beil et al. 2010) packages, respectively. In terms of getting equilibrium points, several packages, such as nleqslv (Hasselman and Hasselman 2018), rootSolve (Soetaert and Soetaert 2009), and BB (Varadhan and Gilbert 2010), are helpful to solve systems of nonlinear equations.

Although the packages offer a wide range of techniques for solving dynamical systems (Soetaert et al. 2012), they suffer from inadequate integration with other tools and frameworks, such as data preprocessing and visualisation libraries. Graphical methods, such as sketching phase velocity vector fields and specific phase curves, are valuable in analysing the potential behaviour of a population under different initial conditions (Percival and Richards 1982; Aron and May 1982; Traulsen et al. 2004). Analytical techniques for investigating a dynamical system involve the identification of equilibrium points and subsequent stability analysis (Begon and Bowers 1995; Murase et al. 2005; Roy and Holt 2008; Saha and Samanta 2020; Azoz and Hussien 2021). These techniques enable researchers to uncover general properties of complicated systems such as pest-pathogen dynamics controlled by pesticides (Saha and Samanta 2020), interactions between predation and infectious disease (Roy and Holt 2008), and the effects of multiple species on insect outbreaks (Dwyer et al. 2004). In practice, however, different techniques are often segregated into multiple packages, each with its own documentation and limited interoperability. This fragmented nature of the available tools poses challenges when it comes to effectively utilising differential equations (Uddin and Robillard 2015).

Another issue that arises in ecological management with dynamical systems is the lack of comprehensive guidelines or frameworks. While general models provide a coarse-scale understanding of fundamental ecological processes such as reproduction, competition, predation, and infections (Hanski et al. 2001; Kim and Reinschmidt 2006; Hartskeerl et al. 2011; Lou and Munther 2012; Grandjean et al. 2020; Luo et al. 2022), practical applications encounter complexities in system dynamics due to heterogeneous populations (Dwyer et al. 1997; Brauer and Watmough 2009; Hou et al. 2021) and various forms of nonlinearity (Faust et al. 2015; Cenci and Saavedra 2018). These complexities often involve context-dependent factors, such as age differences in infections (Cattadori et al. 2008; Hou et al. 2021) and predation (Hastings 1983; Yang and Wang 2020), as well as multiple biophysical processes like tree growth (Pan and Paynal 1995; Zavala et al. 2007; Gayler et al. 2008), seed production (Levin et al. 1989; Ye and Sakai 2016; Detto et al. 2022), and photosynthesis (Friston 2002; Hikosaka 2005; Turschwell et al. 2022). These complexities necessitate the development of location-specific models and the examination of model behaviours in specific scenarios, employing appropriate procedures and a range of analytical and numerical techniques. However, the fragmented nature of functionalities across different packages makes it challenging to navigate through such a procedure. The lack of integration and standardised tools hinders their ability to effectively utilise and apply dynamical systems analysis in conservation management.

This paper introduces a new R package ecode, short for ‘ordinary differential equations in ecology’. The primary goal of ecode is to provide a comprehensive and user-friendly integration of differential techniques for the analysis of ordinary differential equation systems in the context of ecological management. By consolidating various methods and functionalities into a single package, ecode seeks to simplify the process of applying differential equations in ecological studies, provide a general procedure of analysing differential equations, and empower researchers and conservationists to gain deeper insights into complex ecological dynamics and make informed decisions for effective conservation management.

## Methods and features

Package ecode offers a comprehensive framework for constructing and analysing ordinary differential equation systems by integrating various graphical, analytical, and numerical techniques. The core concept of building a differential equation system using the ecode package follows a ‘three-cycle’ procedure (see Fig. 1).

**Figure 1.**
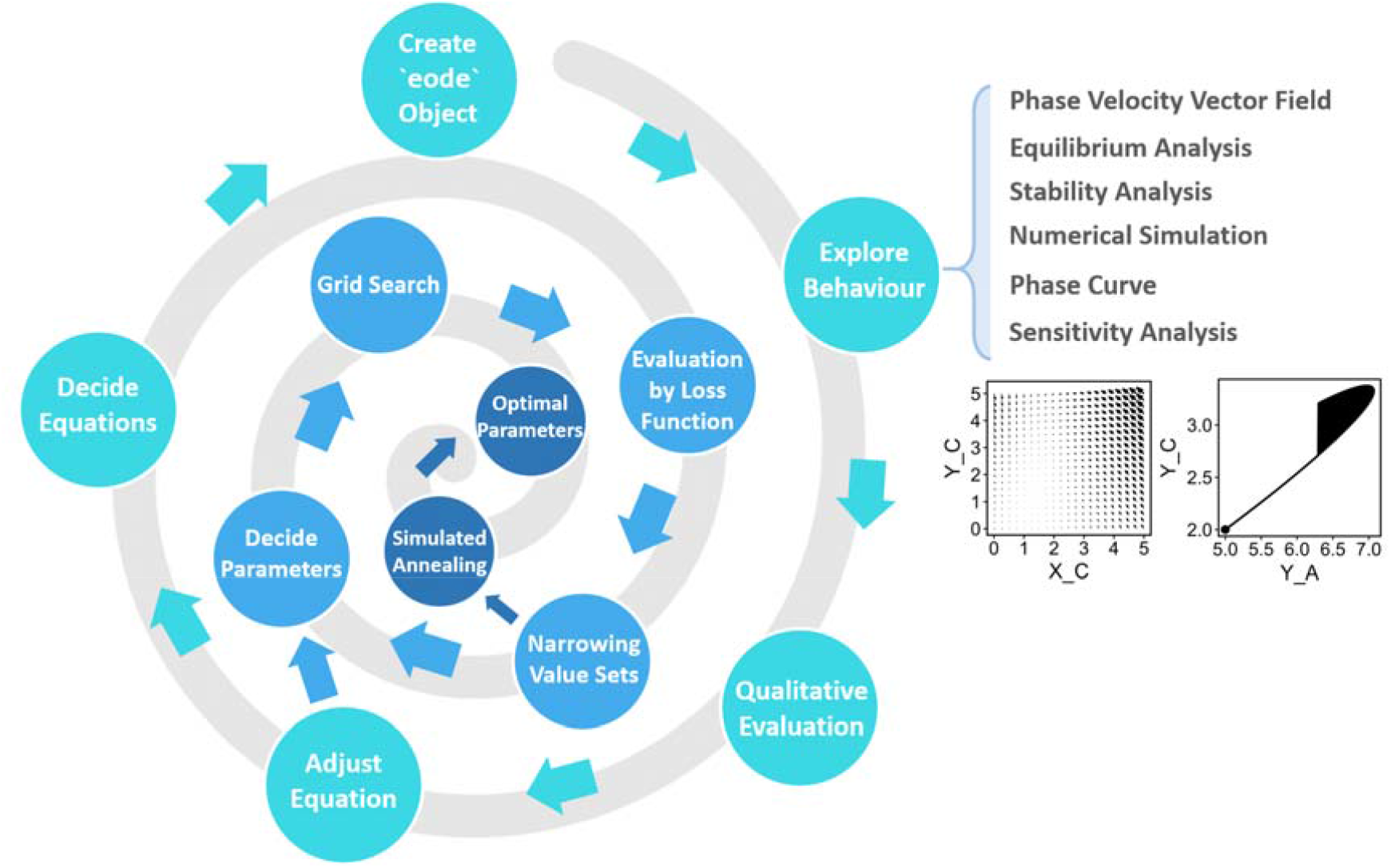
Conceptual diagram illustrating the general procedure for constructing and analysing ordinary differential equations using the ecode package. The construction of a model involves three adaptive cycles. In Cycle 1 (outer side of the spiral), the main goal is to identify an appropriate mathematical formulation and explore a wide range of parameter values that produce qualitative alignment with real-world observations. This stage involves the utilisation of graphical, analytical, and numerical toolkits to analyse the behaviour of the model. During this phase, processes that exhibit scales that are either too small or too large should be removed from the model. Moving on to Cycle 2 (middle of the spiral), users will employ grid search methods to systematically narrow down the parameter space. Finally, in Cycle 3 (inner side of the spiral), users will employ simulated annealing methods with training data to obtain optimal parameter values.

In Cycle 1, users initiate the process by formulating equations based on prior knowledge, creating an ‘eode’ object that represents a differential equation system using the eode() function. Once the object is created, users can employ various techniques to explore the model’s behaviour. They can start by sketching the phase velocity vector field using the plot() function. Equilibrium and stability points can be analysed using functions such as eode_get_cripoi(), eode_is_stapoi(), eode_is_stanod(), eode_is_stafoc(), eode_is_unsnod(), eode_is_unsfoc(), eode_is_saddle(), and eode_is_centre(). In addition to analytical methods, numerical simulations can be performed with different parameter sets using functions eode_proj() and eode_sensitivity_proj(). By conducting simulations and sketching phase curves using plot() and pc_plot() functions, users can gain insights into how the ecological system responds under various conditions. This stage is crucial as it ensures that the model generates patterns that align with observations in real ecological communities (Grimm and Railsback 2005; Brown et al. 2011; Hjøllo et al. 2021). Terms of inappropriate scale may have little impact on the predictive outcomes and should be excluded (Rinaldi and Scheffer 2000; Wu and David 2002; Pásztor et al. 2016).

During Cycle 1 (outer side of the spiral in Fig. 1), it is highly recommended for users to carefully determine a broad range of parameter values that allow the models to generate qualitatively reliable predictions. In epidemiology, for instance, even a slight change in behaviour can have a significant impact on slowing disease progression (Del Valle et al. 2005; Atkeson 2021), ultimately reducing the burden of a pandemic (Ngonghala et al. 2020). To ensure accurate predictions, it is crucial to establish the appropriate interval of values for key parameters such as contact patterns (d’Onofrio and Manfredi 2009) and mortality rate (Jeger et al. 2004). By carefully selecting these parameter values within a realistic range, users can improve the reliability and validity of their models, enabling them to capture the essential dynamics of the ecological system under consideration.

Once a broad parameter space is determined, the procedure transitions to Cycle 2 (middle of the spiral in Fig. 1), where users initiate quantitative assessments to refine the parameter space. The basic idea is to evaluate the deviations between model predictions and real observations using a quadratic loss function. To facilitate this process, the ecode package offers the eode_gridSearch() function, which performs grid search methods to calculate the loss function values for each parameter combination. This aids in narrowing down the search for optimal parameters by identifying a range with relatively small loss function values.

Finally, in Cycle 3 (the inner side of the spiral in Fig. 1), all the necessary preparations have been made to determine the optimal values for the parameter set. The ecode package offers the eode_simuAnnealing() function, which utilises a simulated annealing algorithm to identify the parameter values that yield the minimum loss function value. The simulated annealing algorithm operates by perturbing the parameters at initially high temperatures and gradually cooling down the system. During each annealing step, if the loss function value is smaller than the value in the previous step, the parameter set is updated. Once the optimal parameter values are determined, users can obtain the final model and proceed to explore its behaviour, make predictions, and generate management suggestions. With the optimised parameter set, the model becomes a valuable tool for analysing ecological communities, providing insights into their dynamics and supporting decision-making processes.

Flexible and easy-to-read grammar is an essential aspect of user-friendly software packages, as demonstrated by popular tools such as ggplot2 (Wickham 2011), plotly (Sievert et al. 2023), and dplyr (Wickham et al. 2014). In the realm of differential equation modelling, package ecode distinguishes itself by employing a user-friendly grammar that fosters ease of understanding. The ecode package employs straightforward grammar for creating an ‘eode’ object. When using the eode() function, each argument is a user-defined R function. Model variables are specified as function arguments without default values, while model parameters are defined as function arguments with default values. To ensure that population numbers or abundances remain non-negative, the eode object automatically possesses constraints that confine model variables to positive values. To further customise constraints, users can modify the values of the ‘constraint’ argument in the eode() function using intuitive mathematical expressions such as ‘X<100’. Additionally, even after establishing an eode object, users retain the flexibility to modify parameter settings or constraints using the eode_set_constraint() and eode_set_parameter() functions, allowing for iterative refinement and adjustment of the model.

In addition, the ecode package employs the S3 inheritance mechanism, enabling the use of an intuitive grammar to access printing services and visualisation libraries. For sketching the phase velocity vector field, users can simply employ the plot() function on an eode object.

The system automatically invokes the plot.eode() function, which is specifically defined to generate a two-dimensional plot of the phase velocity vector field by default. When conducting numerical simulations with the eode_proj() function, users obtain an object of the ‘pc’ class, representing a phase curve. To visualise the simulation trajectory, users can apply the plot() function directly on the object of pc class. For sensitivity analysis, the eode_sensitivity_proj() function returns an object of the ‘pcf’ class, representing a phase curve family. Similar to the pc class, users can utilise the plot() function on objects of the pcf class to visualise how changes in initial conditions and parameters influence the model predictions. Moreover, the S3 inheritance mechanism is utilised to overload the ‘print’ function in R for objects of the eode, pc, and pcf classes. This allows for the convenient printing of brief details about the model information and phase curve properties, providing users with informative summaries of their models and analysing results.

Dependencies among R packages also deserve careful considerations (Theußl et al. 2011; Decan et al. 2016; Gu and Hübschmann 2022). While importing external packages reduces code replication, it also introduces risks, as a single dependency failure can render the entire package unusable (Cox 2019). Updates in package dependencies, such as changes in function names, can also lead to unintended consequences and errors that are difficult to trace (Decan et al. 2016; Decan et al. 2017). While integrating various analysis techniques from different R packages is an option, ecode prioritises minimising external dependencies to ensure robustness for practical uses. This approach aims to avoid package failures caused by updates or changes in one or several dependencies. As a result, ecode relies only on core packages in R, namely ggplot2 (Wickham 2011), rlang (Henry et al. 2023), and stringr (Wickham, 2010), minimising the risk of unforeseen issues arising from external dependencies.

## Example

This section demonstrates the practical usage of the ecode package, following the ‘three-cycle’ procedure (Fig. 1). To illustrate its application, we will consider a commonly used ecological system based on the susceptible-infected model (Keeling 2008; Martcheva 2015). The imaginative population are categorised into four classes: susceptible child (*X*_*C*_), infected child (*Y*_*C*_), susceptible adult (*X*_*A*_), and infected adult (*Y*_*A*_). The model incorporates several processes, including growth, natural mortality of the child class, disease transmission, disease-induced mortality, and reproduction.

### Cycle 1: Adapt mathematical formulation and parameter space

To begin, we can create an eode object using the eode() function. Each argument of the eode() function should be an R function representing a single equation that defines the rules for calculating phase velocity vectors. As mentioned earlier, the arguments without default values in the function correspond to model variables, while the arguments with default values represent model parameters.

We first create an eode object using the eode() function. Each argument of the eode() function should be an R function representing a single equation that defines rules for calculating phase velocity vectors. As mentioned earlier, the arguments without default values in the function correspond to model variables, while the arguments with default values represent model parameters. Consider the following system:

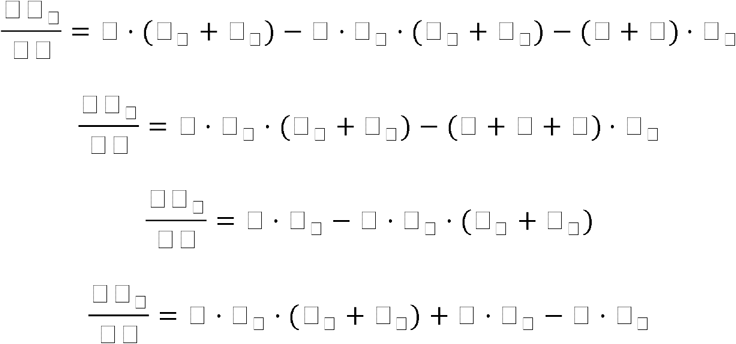

Each of the model variable denotes the population size for each class, and *v* stands for reproduction rate (0.15 in this case), *β* for transmission rate (0.1 in this case), μ for background mortality rate of child class (0.15 in this case), and □ for growth rate (0.04 in this case), *ρ* for the rate of disease-induced mortality (0.2 in this case).

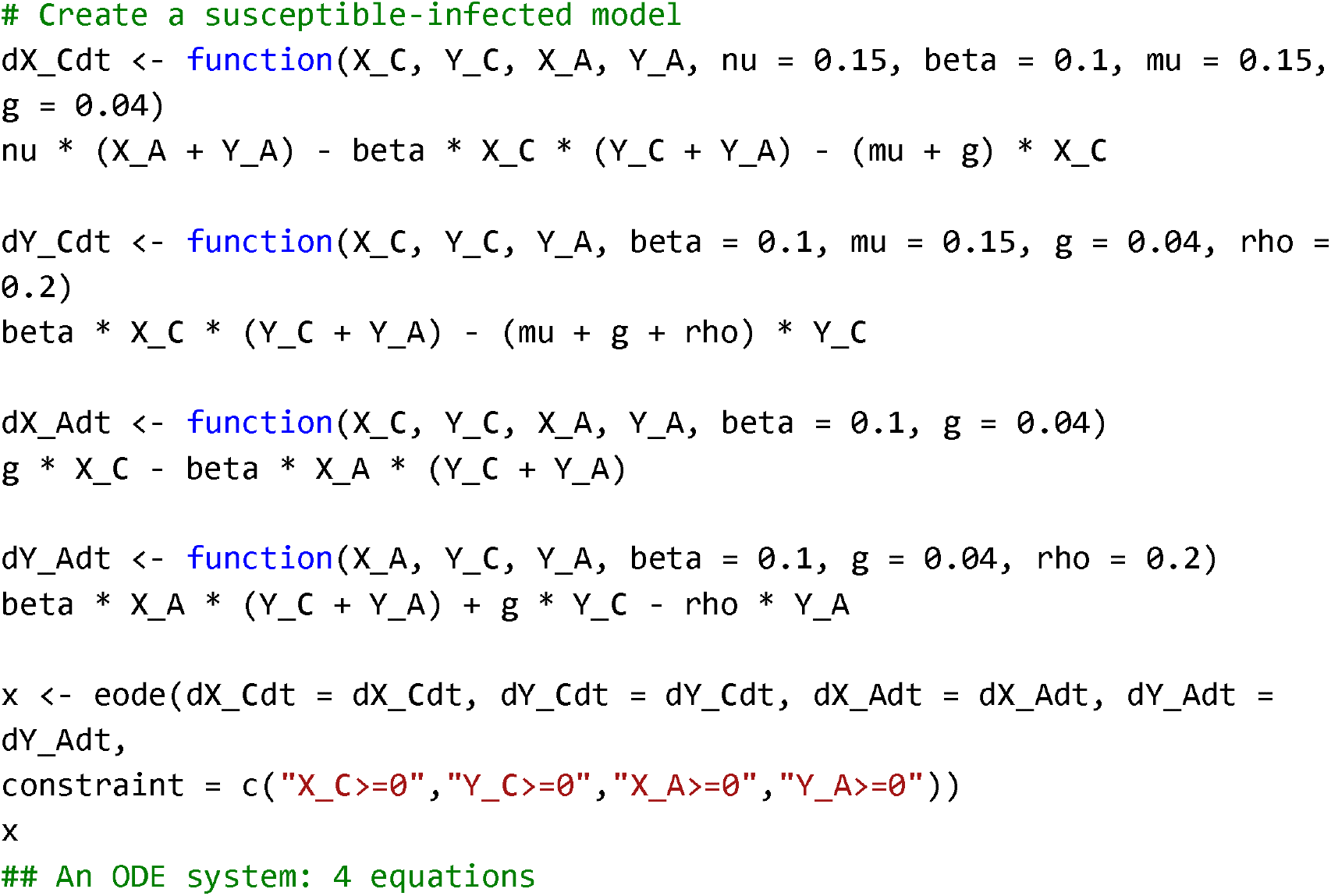
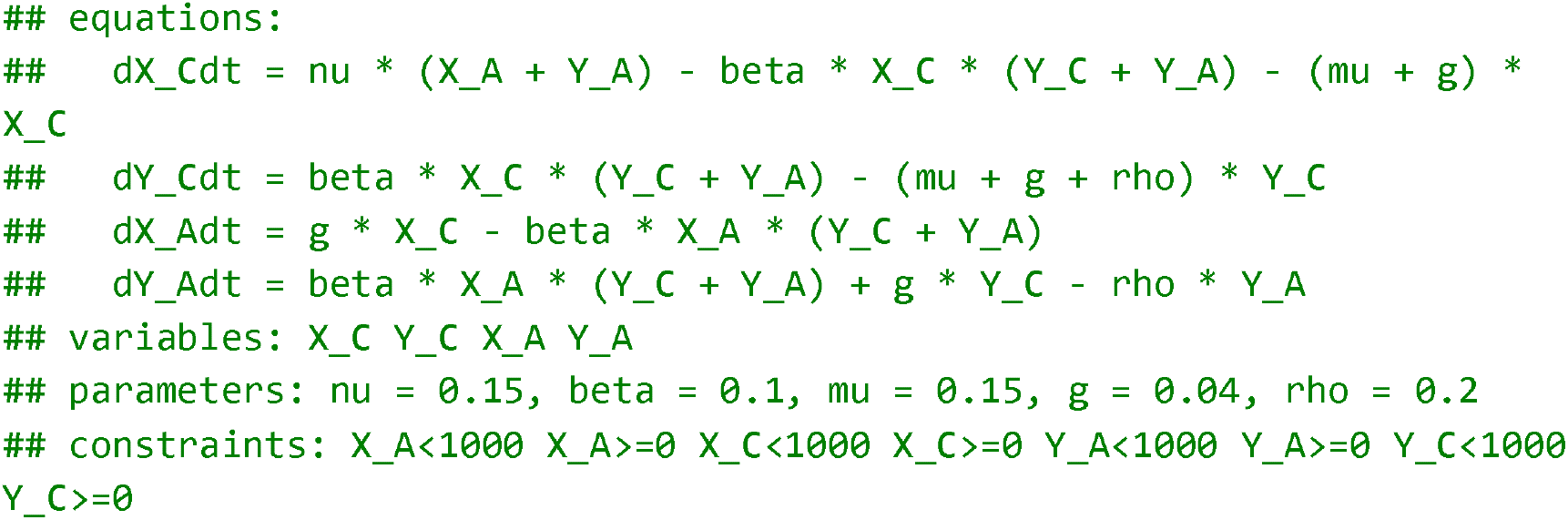

The ‘constraint’ argument of the eode() function allows us to define boundary limits for model variables. By default, the eode() function sets the range for model variables as 0 to 1000. The output of printing an eode object shows details about the mathematical formulations, model variables, parameters and their corresponding values, as well as constraints of model variables.

Then, we explore the phase velocity vector field of the epidemiological system using plot() function (Fig. 2). When the model involves more than two variables, the plot() function automatically assigns the additional variables with their middle values within their boundaries. This approach allows us to visualise the phase velocity vector field in a two-dimensional space. However, it is also allowed to customise the value settings for these extra variables using the ‘set-covar’ argument of the plot() function.

**Figure 2.**
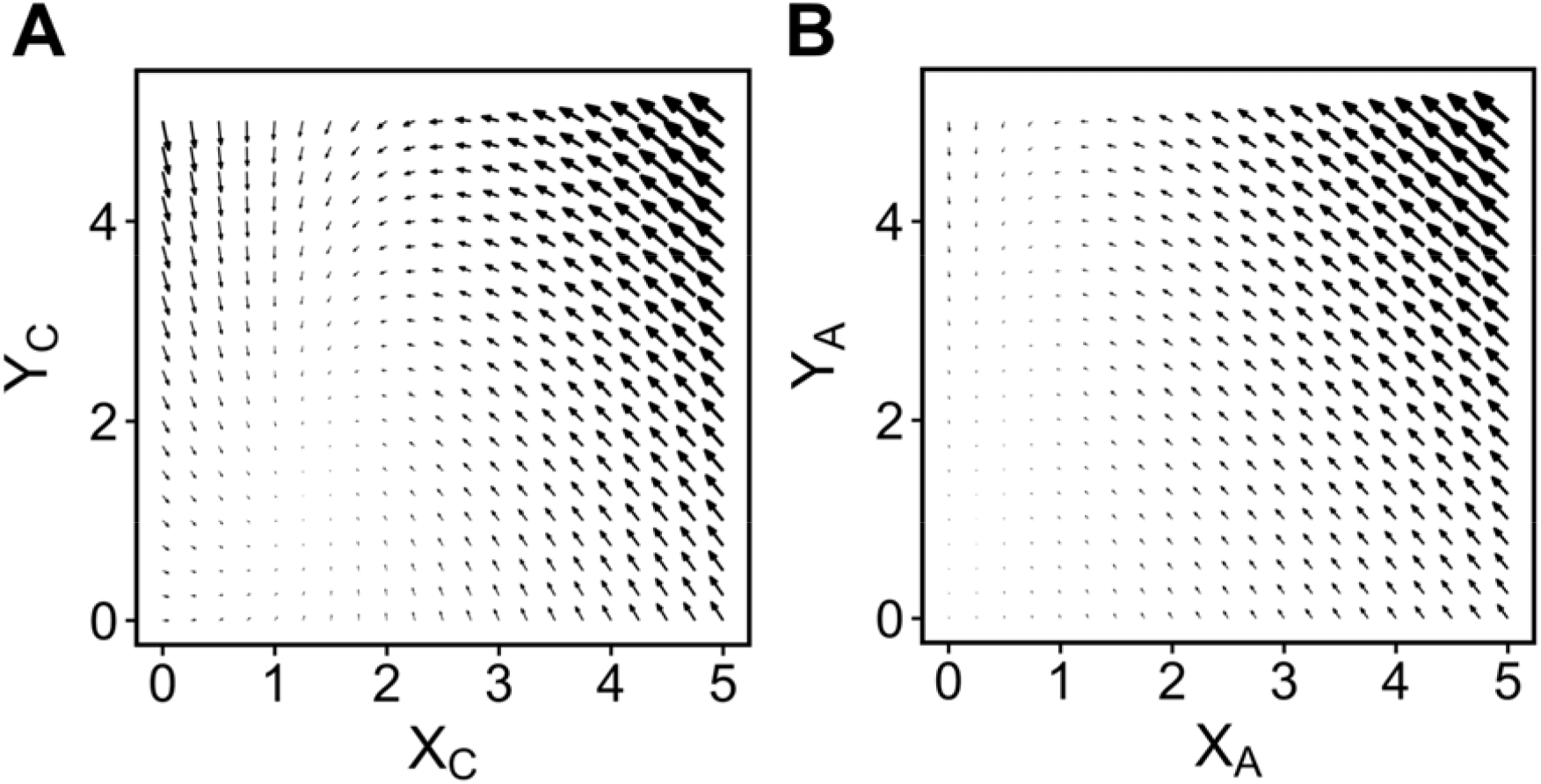
Plot of phase velocity vector field under the condition of (A) *X*_*A*_ = *Y*_*A*_ = 2.5; (B) *X*_*C*_ = *Y*_*C*_ = 2.5, where *X*_*A*_, *Y*_*A*_, *X*_*C*_, and *Y*_*C*_ stand for population size of susceptible adult, infected adult, susceptible child, and infected child classes, respectively. Each arrow indicates the direction of phase velocity in a single phase point. The length and weight of the arrows indicates the magnitude of the phase velocity vector. The plot is created by the plot() function acted on objects of eode class.

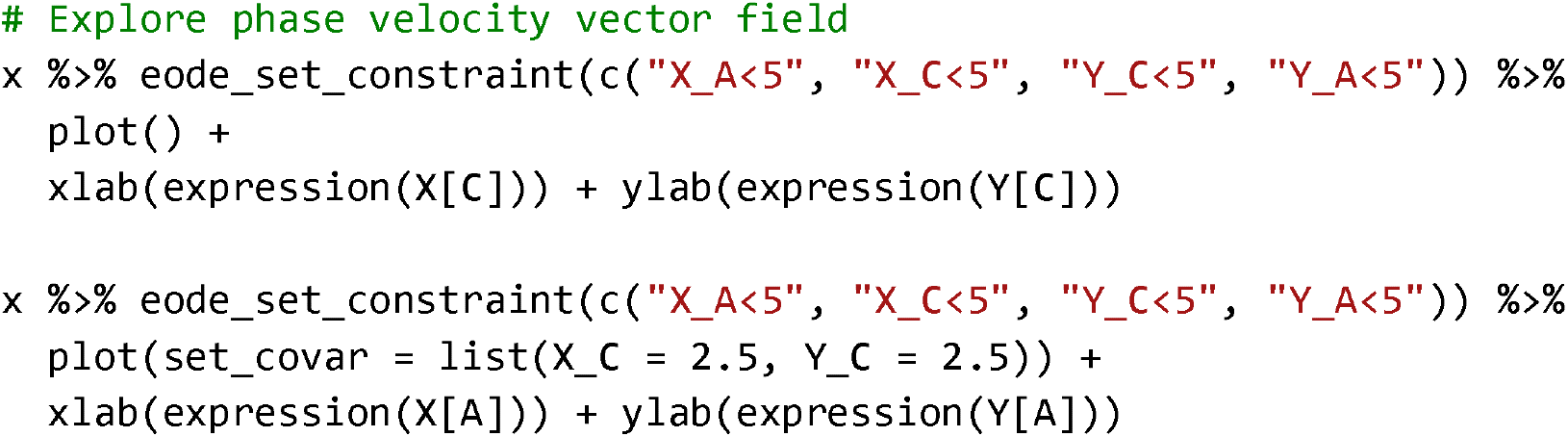

The phase velocity vector field (Fig. 2) suggests the presence of an equilibrium point when all the model variables are close to zero. To identify this equilibrium point, we can utilise the eode_get_cripoi() function, which employs Newton iteration methods to search for the equilibrium point from an initial point (*X*_*C*_, *Y*_*C*_, *X*_*A*_, *Y*_*A*_) = (1, 1, 1, 1).

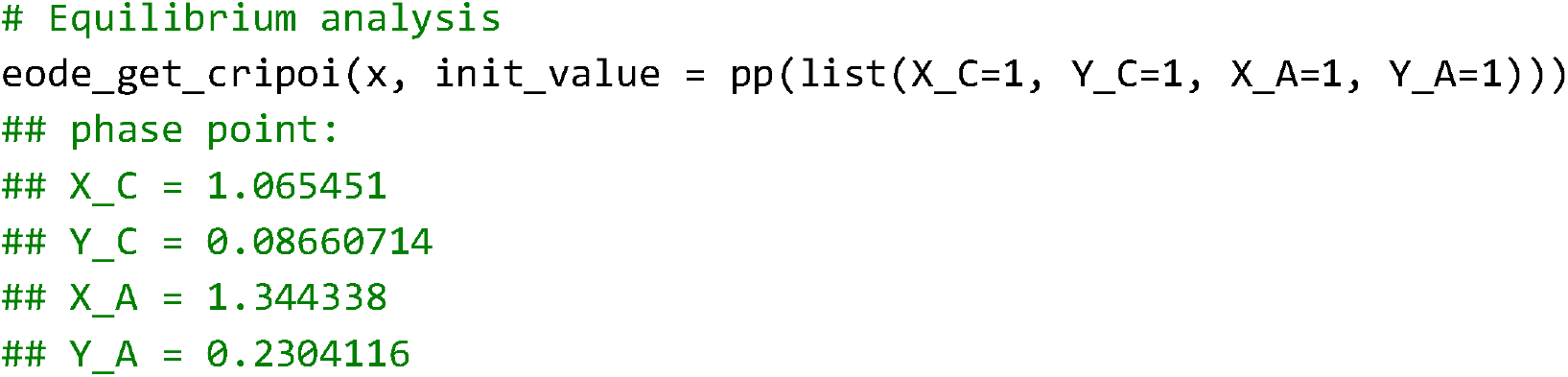

We can further assess whether the equilibrium point is a stable point using the function eode_is_stapoi(). A similar manner applied to the identification of stable nodes, stable foci, unstable nodes, unstable foci, saddles, and centres.

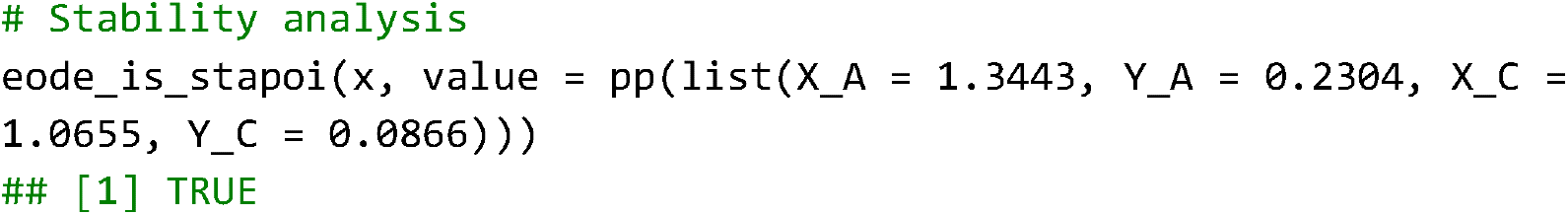
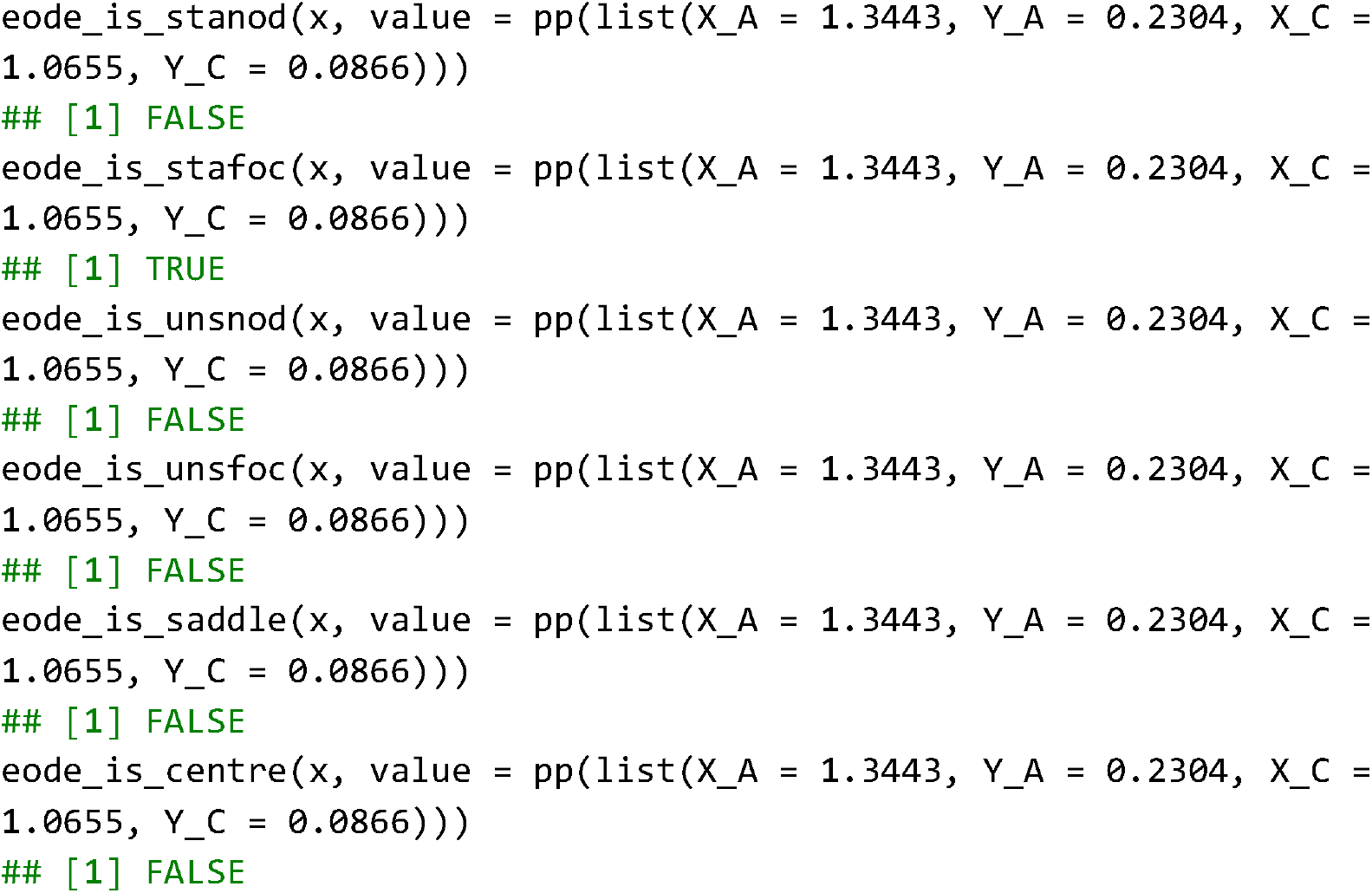

To perform numerical simulations, we can use the eode_proj() function, which returns a pc object representing a phase curve. Function plot() and pc_plot() for pc objects can be used to visualise the model’s dynamics (Fig. 3) and the movement along the phase line (Fig. 4).

**Figure 3.**
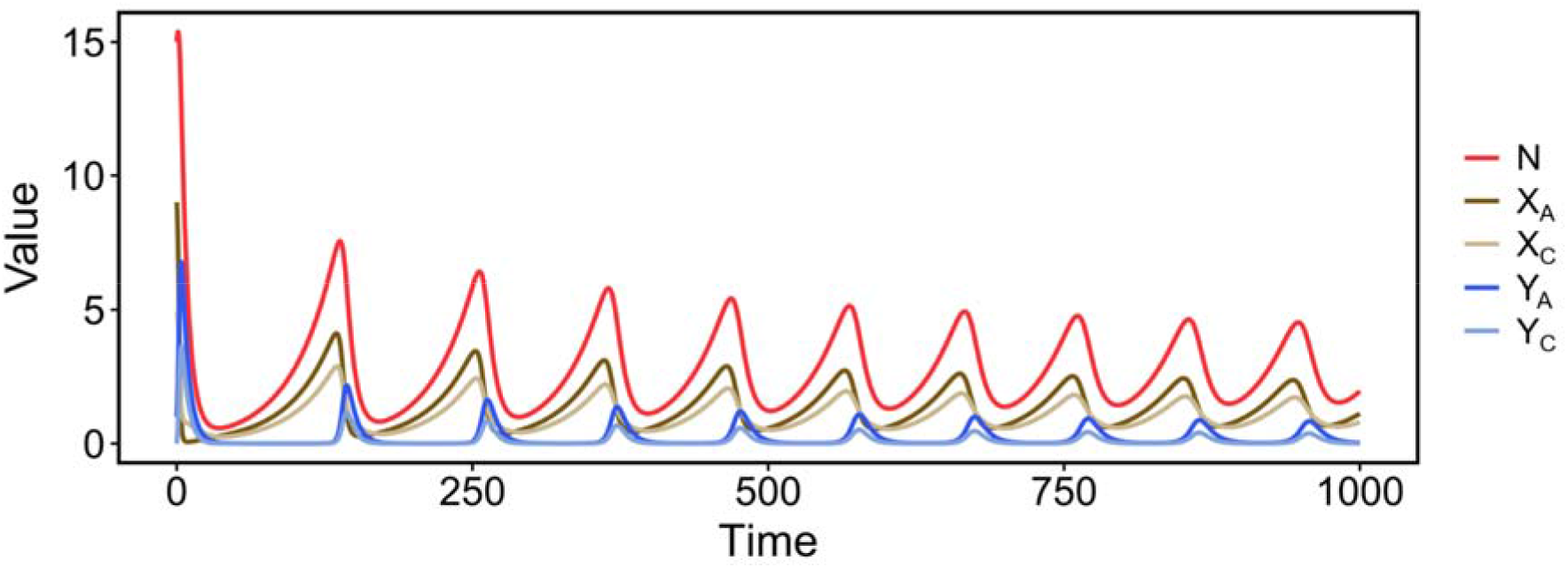
Epidemiological dynamics of the model. The *x*-axis represents the time scale (unit in year) and the *y*-axis represents population size of different classes: *X*_*A*_ for susceptible adult, *X*_*C*_ for susceptible child, *Y*_*A*_ for infected adult, and *Y*_*C*_ for infected child. *N* stands for the total population size. Created by the function plot() for pc objects. Simulation is based on the initial condition of *X*_*A*_(0) = 9, *Y*_*A*_(0) = 1, X_C_(0) = 5, *Y*_*C*_(0) = 0.

**Figure 4.**
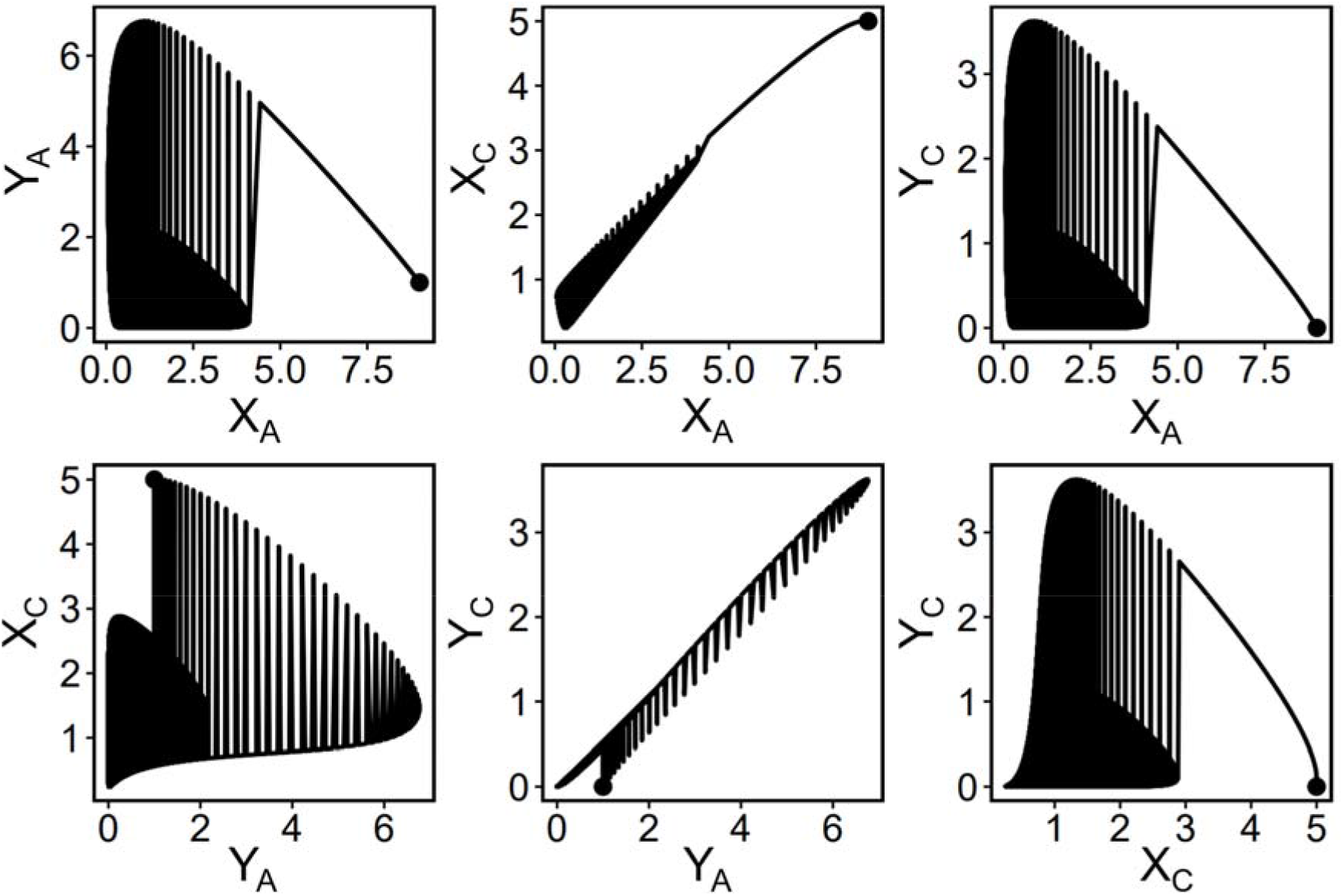
Phase line generated by a single numerical simulation. *X*_*A*_ stands for susceptible adult, *X*_*C*_ for susceptible child, *Y*_*A*_ for infected adult, and *Y*_*C*_ for infected child. Created by the function pc_plot(). Simulation is based on the initial condition of *X*_*A*_(0) = 9, *Y*_*A*_(0) = 1, X_C_(0) = 5, *Y*_*C*_(0) = 0.

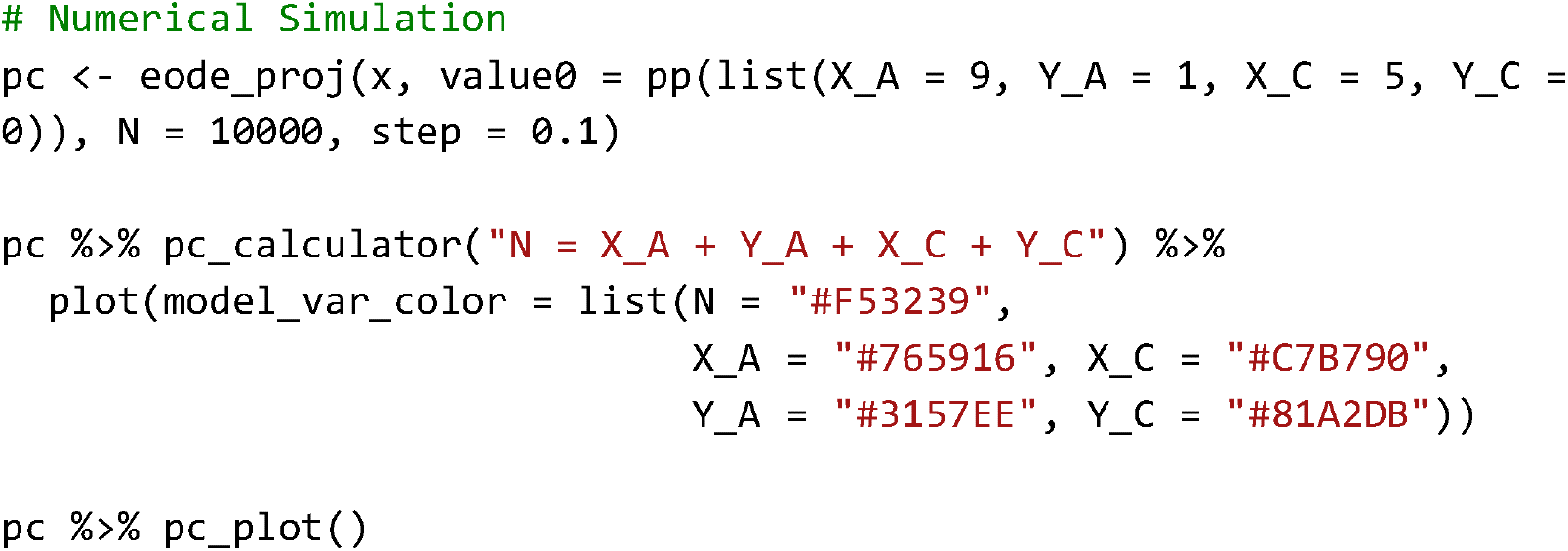

To explore how variations in initial conditions and parameter values influence the model behaviour, we can perform sensitivity analysis using the function eode_sensitivity_proj(), which creates a pcfamily object that can be visualised by the function plot() (Fig. 5). For instance, if we want to examine the effect of changing the parameter β from 0.05 to 0.15 with a step size of 0.05, we can specify ‘beta = c(0.05, 0.1, 0.15)’ in the ‘valueSpace’ argument of the eode_sensitivity_proj() function. A similar manner applied to change in initial conditions.

**Figure 5.**
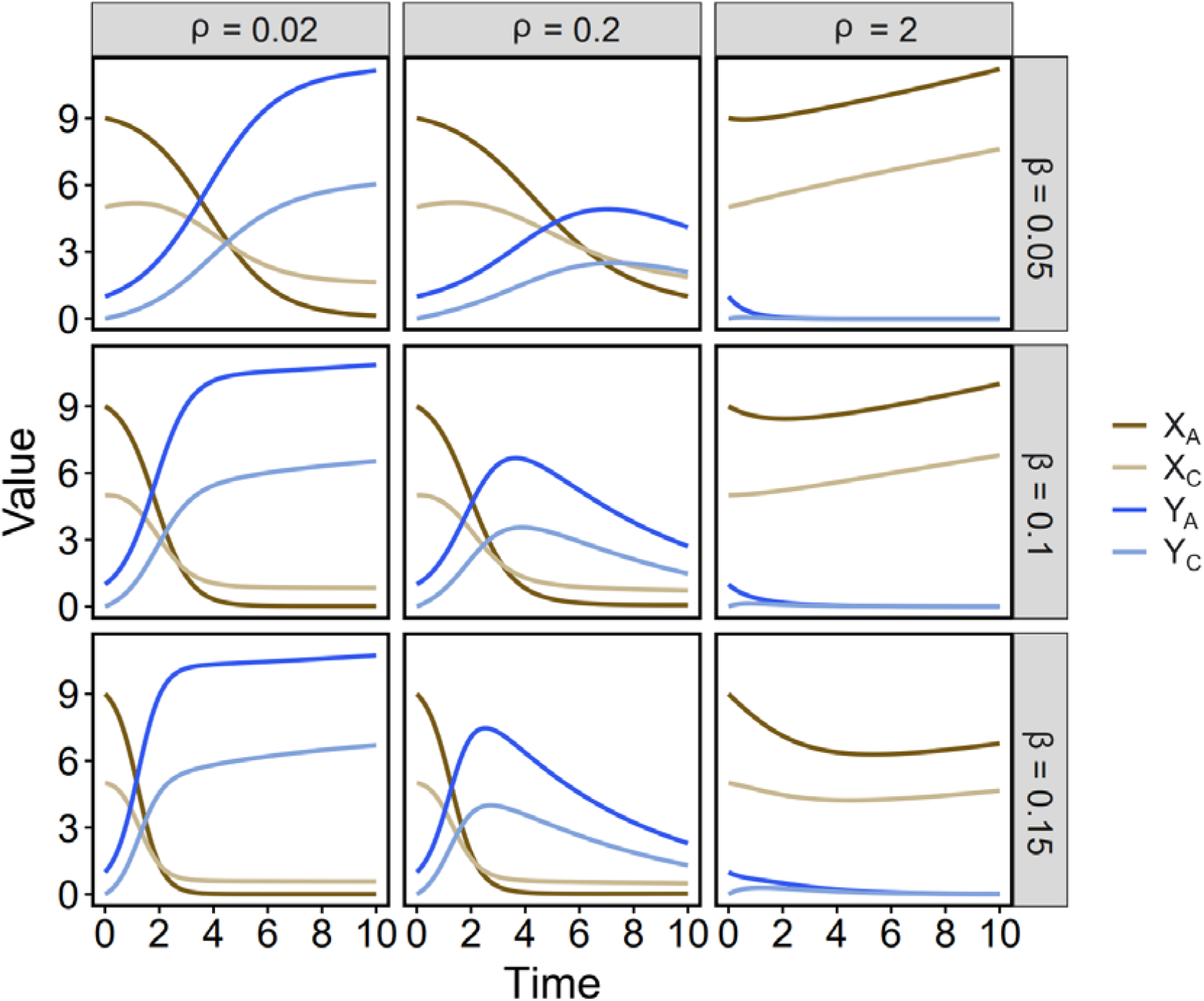
Effects of changes in transmission rate (*β*) and disease-induced mortality rate (*ρ*) on epidemiological dynamics. The *x*-axis represents the time scale (unit in year) and the *y*-axis represents population size of different classes: *X*_*A*_ for susceptible adult, *X*_*C*_ for susceptible child, *Y*_*A*_ for infected adult, and *Y*_*C*_ for infected child. Created by the function plot() for pc objects. Simulation is based on the initial condition of *X*_*A*_(0) = 9, *Y*_*A*_(0) = 1, X_C_(0) = 5, *Y*_*C*_(0) = 0.

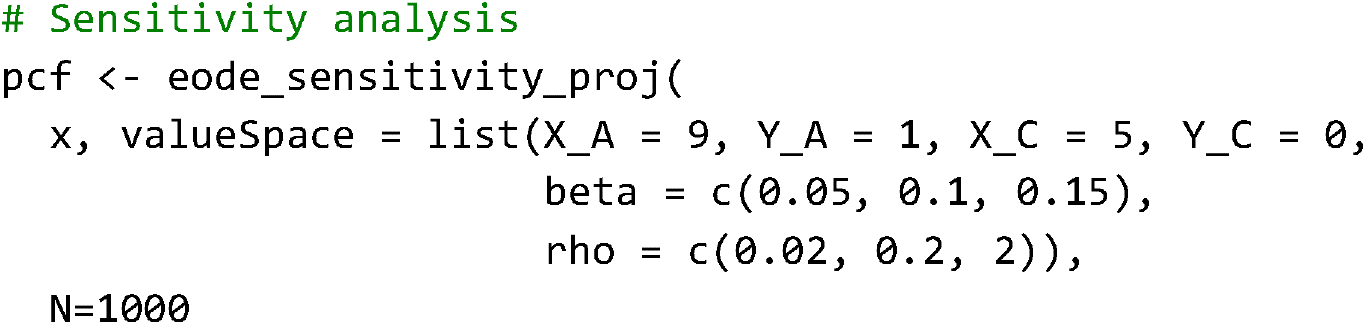
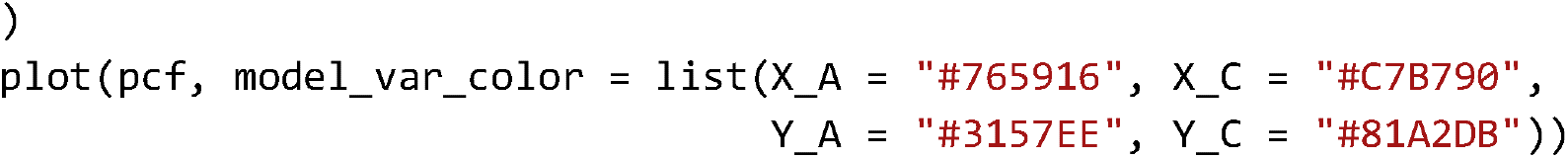

### Cycle 2: Narrow down parameter space

To perform a grid search method, we need to define training data using the pdata() function. Assuming we have *n* observations in the field, then the ‘init’ argument of the pdata() function should be a data frame with *n* rows, each row representing a single observation with specific initial value (e.g. number of susceptible individuals at the beginning year). Argument ‘*t*’ should be a vector of length *n*, indicating the time of observation relative to the initial year.

Argument ‘lambda’ should be a data frame of *n* rows, each row representing a single observation with specific values of observed variables (in this case, the incident rate of infectious disease). The last argument ‘formula’ defines how to calculate the observed variables.

Once the training data is defined, we can invoke the eode_gridSearch() function and find parameters where loss function values are minimised, thus narrowing down the potential parameter space. In this case, we vary the transmission rate (β) from 0.05 to 0.15 with a step of 0.01 and find that the minimal loss function value is achieved when β = 0.11.

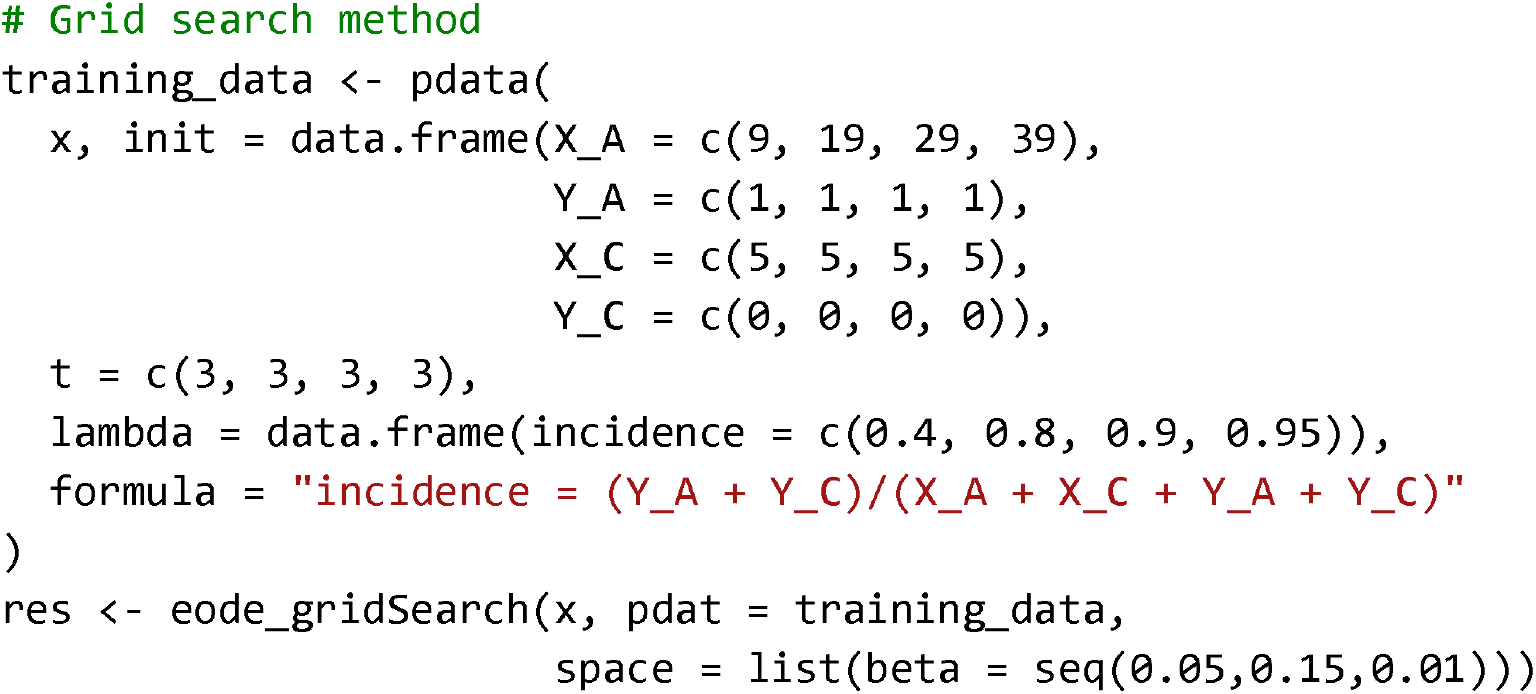

#### Cycle 3: Optimise parameter settings

Finally, to determine the optimal parameter settings, we can utilise a simulated annealing algorithm. In this case, since we have identified that a minimal loss function value is achieved when β = 0.11 through grid search methods, we can set β = 0.11 using the eode_set_parameter() function. Subsequently, we can invoke the eode_simuAnnealing() function to initiate the simulated annealing process and obtain the most suitable value of transmission rate β that aligns with the observed data, making sure that the model provides accurate predictions.

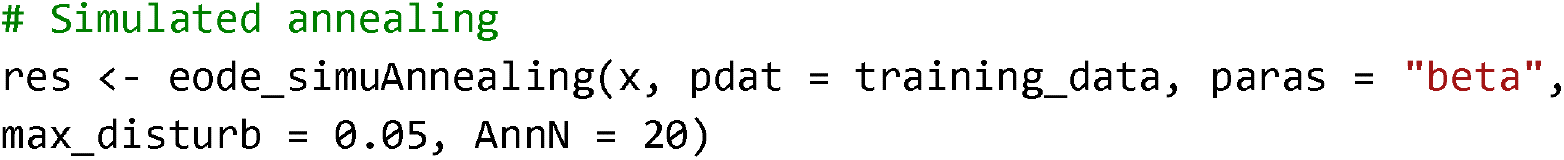

## Conclusion

In conclusion, the ecode package offers a comprehensive framework for building and analysing ordinary differential equation systems in the study of population of community ecology as well as biodiversity management. By following the three-cycle procedure, users can confidently develop and refine their models with a wide range of graphical, analytical, and numerical techniques. The package features intuitive grammar and user-friendly interface, enhancing accessibility and allowing researchers to easily explore and understand model behaviour. The incorporation of grid search methods and simulated annealing empowers users to iteratively optimise their model parameters using observed data. With minimal external dependencies, ecode ensures stability and reliability, making it a valuable tool for ecological modelling and management practices. By leveraging the capabilities of the ecode package, researchers and conservationists can gain insights into complex ecological systems, make informed predictions, and generate valuable recommendations for effective environmental management.

## Data availability statement

Package ecode is available on GitHub at https://github.com/HaoranPopEvo/ecode.

